# Feed-forward inhibition fine-tunes response timing in auditory-vocal interactions

**DOI:** 10.1101/2021.09.03.458890

**Authors:** Philipp Norton, Jonathan Benichov, Margarida Pexirra, Susanne Schreiber, Daniela Vallentin

## Abstract

The ability to regulate vocal timing is a fundamental aspect of communicative interactions for many species, including conversational speech among humans, yet little is known about the neural circuitry that regulates the input-dependent timing of vocal replies. Exploring this topic in the zebra finch premotor area HVC, we identify feed-forward inhibition as a key regulator of vocal response timing. Based on a spiking network model informed by behavioral and electrophysiological data from communicating zebra finches, we predicted that two different patterns of inhibition regulate vocal-motor responses. In one scenario, the strength of production-related premotor inhibition translates into plasticity in vocal response delays. In the other scenario, fast transient interneuron activity in response to auditory input results in the suppression of call production while a call is heard, thereby reducing acoustic overlap between callers. Extracellular recordings in HVC during the listening phase confirm the presence of auditory-evoked response patterns in putative inhibitory interneurons, along with corresponding signatures of auditory-evoked activity suppression. The proposed model provides a parsimonious framework to explain how auditory-vocal transformations can give rise to vocal turn-taking and highlights multiple roles of local inhibition for behavioral modulation at different time scales.

## Introduction

### Behavioral Importance of Vocal turn-taking

A defining characteristic of spoken conversations is the alternating exchange of vocalizations, often with rapid transitions between speakers and minimal overlap of speech (Levinson, 2016). This example of vocal turn-taking requires precise control of the onsets of vocalizations, with individual speakers typically responding to their conversational partners within ~250 ms, although average speeds can vary across linguistic cultures (Stivers et al., 2009).

The ability to coordinate vocalizations in an interspersed manner precedes spoken language developmentally and evolutionarily, extending to other species ranging from non-human primates to birds and frogs (Pika et al., 2018). In all cases, vocal interactions generally require perceiving relevant acoustic signals and initiating exact motor commands to generate an appropriate vocal reply. In the case of vocal turn-taking, each interlocutor delays or withholds a response while listening to the other. This social form of sensorimotor coordination reduces acoustic overlap, thereby maintaining unmasked signal transmission and detection. Although this behavior is wide-spread, little is known about how brain circuits flexibly control whether and when to respond to a partner’s vocalizations.

### Forebrain control of coordinated vocal timing in zebra finches

The zebra finch has served as a tractable model system for studying the neuroethology of developmental vocal learning (Immelmann, 1968; Scharff & Nottebohm, 1991; Zann, 1996; Tchernichovski et al., 2001). Due to their distributed nucleated brain architecture (Nottebohm et al., 1976) songbirds are particularly well suited to study the dedicated neural circuits underlying vocal learning and production (Hahnloser et al., 2002; Brainard & Doupe, 2002; Long et al., 2010; Okubo et al., 2015; Vallentin et al., 2016), The vocal-motor pathway has been studied extensively to understand the neural mechanisms underlying production of courtship song, which male zebra finches perform in a uni-directional rather than turn-taking manner. Recently, the convergence of behavioural, anatomical, and electrophysiological evidence has indicated that the zebra finch forebrain “song system” is not solely dedicated to the learned performance of complex courtship song, but that the descending forebrain vocal-motor pathway is also involved in the production of acoustically simpler innate affiliative calls (Hahnloser et al., 2002; Ter Maat et al., 2014; Benichov et al., 2016; Shaughnessy et al., 2019; Benichov & Vallentin, 2020; Ma et al., 2020).

Zebra finches engage in pair-specific antiphonal exchanges of short calls, often coordinating calls with one-another within the context of a larger group (Gill et al., 2015; Ter Maat et al., 2014; Elie & Theunissen, 2020). This example of vocal turn-taking requires precise regulation of call timing relative to the calls of others. In controlled settings, birds can be driven to adapt their call timing to avoid “jamming” (i.e. overlapping with) the calls of another bird or temporally predictable call playbacks (Benichov et al., 2016; Benichov & Vallentin, 2020). Blocking the influence of the forebrain vocal-motor pathway by lesioning the song system output nucleus RA (Robust nucleus of the Arcopallium) or through pharmacological inactivation of the directly upstream premotor nucleus HVC (proper name) drastically impairs the temporal precision of the call response and consequently, jamming avoidance.

Electrophysiological recordings within the vocal-motor pathway of awake-behaving birds have identified bursts of activity in HVC premotor neurons related to call onsets (Hahnloser et al., 2002; Ter Maat et al., 2014; Benichov & Vallentin, 2020; Ma et al., 2020). Results from intracellular recordings have implicated the inhibitory activity of HVC interneurons in modulating the sparse bursting of premotor projection neurons that appear to trigger call production. Furthermore, pharmacological manipulation of local inhibition within HVC has profound effects on calling behavior, with disinhibition resulting in significantly faster call response latencies (Benichov & Vallentin, 2020). Here we utilize these previously observed data along with new extracellular recordings in awake birds listening to call playbacks to provide the empirical basis for a mathematical model of a vocal timing control circuit.

### Modelling a vocal timing control circuit

While the previous experimental results (Ma et al., 2020; Benichov et al., 2016) imply the involvement of HVC in controlling the timing of calls in vocal interactions, the exact functional interplay between identified cell types within this circuitry is unknown. In this study we developed a leaky integrate-and-fire (LIF) neuron-based spiking network model composed of HVC premotor and interneurons as well as upstream auditory neurons and evaluated the plausibility of connectivity profiles and circuit mechanisms in terms of their consistency with experimental observations.

The proposed mathematical model of HVC’s involvement in call perception and timing allowed us to systematically explore multiple components of this vocal circuit: 1) The interplay between excitatory drive and local inhibition; and 2) the interactions between sensory input during listening and premotor output that leads to a vocalization. The interpretation of experimental data can be limited by the need to align and analyze activity in relation to either an incoming auditory stimulus or the vocalization, potentially obscuring potential interactions between the two. This model provides a more flexible framework, enabling the direct simulation of experimentally less tractable conditions including circuit connectivity, helping us to dissect the roles of specific circuit components in the control of vocal response timing. Specifically, the generation of multiple scenarios in which premotor activity occurs at different time points relative to an arriving auditory stimulus enabled us to derive a plausible mechanism for how inhibition regulates call onset times that proved consistent with subsequent experimental test based on the model’s predictions.

## Results

### A spiking network model for call production-related activity in HVC

We developed a spiking network model consisting of leaky integrate-and-fire neurons connected through bi-exponential current-based synapses (Roth & van Rossum, 2010; p. 143) with the initial aim of accurately replicating the call-related activity of HVC premotor neurons and interneurons (Benichov & Vallentin, 2020) on a microcircuit level. Compared to more biophysically realistic Hodgkin-Huxley type neuron models, LIF models have fewer parameters and are more computationally efficient in numerical simulations. Integrate-and-fire neurons have previously been successfully applied in modeling of HVC activity during song production (Li & Greenside, 2006; Cannon et al., 2015; Hamaguchi et al., 2016). Here, the intrinsic neuronal properties, as well as synaptic weights and time constants, were fit to data from electrophysiological studies of zebra finch HVC (Table 1 & 2; Mooney & Prather, 2005; Kosche et al., 2015; Hamaguchi et al., 2016).

**Table 1 -.** Neuron and synapse model parameters. Parameters used for all excitatory (vocal- and auditory-related input, premotor) and inhibitory neurons (interneurons). E_L_: leakage or resting membrane potential, v_reset_: reset potential, v_thresh_: spiking threshold potential, T_m_: membrane time constant, R_m_: membrane resistance, T_decay_e/T_decay_i: decay time constants for excitatory and inhibitory synaptic currents, respectively, T_rise_e/T_rise_i: rise time constants for excitatory and inhibitory synaptic currents, respectively, t_ref_: absolute refractory period.

| | $E_L$<br>[mV] | $V_{reset}$<br>[mV] | $V_{thresh}$<br>[mV] | $\tau_m$<br>[ms] | $R_m$<br>[MΩ] | $\tau_{decaye}$<br>[ms] | $\tau_{risee}$<br>[ms] | $\tau_{decayi}$<br>[ms] | $\tau_{risei}$<br>[ms] | $t_{ref}$<br>[ms] |
| --- | --- | --- | --- | --- | --- | --- | --- | --- | --- | --- |
| **Excitatory** | -75 | -50 | -40 | 16 | 200 | 1.6 | 0.4 | 2.2 | 0.4 | 1 |
| **Inhibitory** | -60 | -70 | -45 | 8 | 200 | 0.6 | 0.5 | 0.6 | 0.5 | 1 |

**Table 2 -.** Network model parameters. Parameters used for the different network models presented in Figures 1, 2 and 5. Capital letters denote the different neuron populations. V: Vocal-related input, I_v_, I_a_: Interneuron, P: Premotor, A: Auditory-related input, T: Tonically active interneuron. The synaptic weight from the vocal-related input population to the silent premotor neuron (yellow in Figure 1B) was 8 pA instead of 20 pA. Parameters that are identical for all models are shaded in gray. Connection probabilities (Conn. prob.) and synaptic weights and delays between populations A, I_a_ and P in the full model are the same as between V, I_v_ and P, except where stated otherwise.

| Model | No Inhibition<br>(Figure 2A) | Tonic Inhibition<br>(Figure 2C) | FF-Inhibition<br>(Figure 1 & 2E) | Full model<br>(Figure 5A) |
| --- | --- | --- | --- | --- |
| Nr. of neurons V | 150 | 150 | 150 | 150 |
| Nr. of neurons I<sub>v</sub> | 30 | 30 | 30 | 30 |
| Nr. of neurons P | 1 | 1 | 1 | 1 |
| Nr. of neurons A | — | — | — | 150 |
| Nr. of neurons I<sub>a</sub> | — | — | — | 25 |
| Nr. of neurons T | — | 120 | — | — |
| Conn. prob. M → I<sub>v</sub> | 0.3 | 0.3 | 0.3 | 0.3 |
| Conn. prob. I<sub>v</sub> → P | — | — | 1 | 1 |
| Conn. prob. V → P | 0.4 | 1 | 1 | 1 |
| Conn. prob. T → P | — | 1 | — | — |
| Syn. weights V → I<sub>v</sub> | 40 pA | 40 pA | 40 pA | 40 pA |
| Syn. weights I<sub>v</sub> → P | — | — | -19 pA | -21 pA |
| Syn. weights V → P | 20 pA | 20 pA | 20 pA | 20 pA |
| Syn. weights T → P | — | -23 pA | — | — |
| Syn. weights A → P | — | — | — | 6 pA |
| Syn. delay V → I<sub>v</sub> | 0.5 ms | 0.5 ms | 0.5 ms | 0.5 ms |
| Syn. delay I<sub>v</sub> → P | 0.4 ms | 0.4 ms | 0.4 ms | 0.4 ms |
| Syn. delay V → P | 0.9 ms | 0.9 ms | 0.9 ms | 0.9 ms |
| Syn. delay T → P | — | 0.4 ms | — | — |

Intracellular recordings of identified RA-projecting premotor neurons in HVC (HVC_(RA)_; Benichov & Vallentin, 2020; henceforth referred to as ‘premotor neurons’) have revealed that they either exhibit a burst of action potentials (2.4 ± 1.2 spikes per burst, mean ± std; average burst onset: −45 to 33 ms relative to call onset) or are hyperpolarized (onset of hyperpolarization: −52 ± 14 ms) shortly before the onset of a produced call (Figure 1A). The model simulated the activity of a representative cell from the set of call-bursting premotor neurons and from a set of premotor neurons that do not burst during calling (“silent” with respect to calls) but are hyperpolarized prior to call onsets (Figure 1B). The activity profile of HVC premotor neurons was modulated by local inhibitory interneurons within HVC (Kosche et al., 2015; Markowitz et al., 2015). During calling, a subset of these interneurons transiently increased its firing rate prior to call-related premotor bursts, also coinciding with the onset of hyperpolarization in the silent premotor neurons (Figure 1A). The model reproduced this firing rate increase and timing relative to call production (Figure 1B).

**Figure 1 –.**
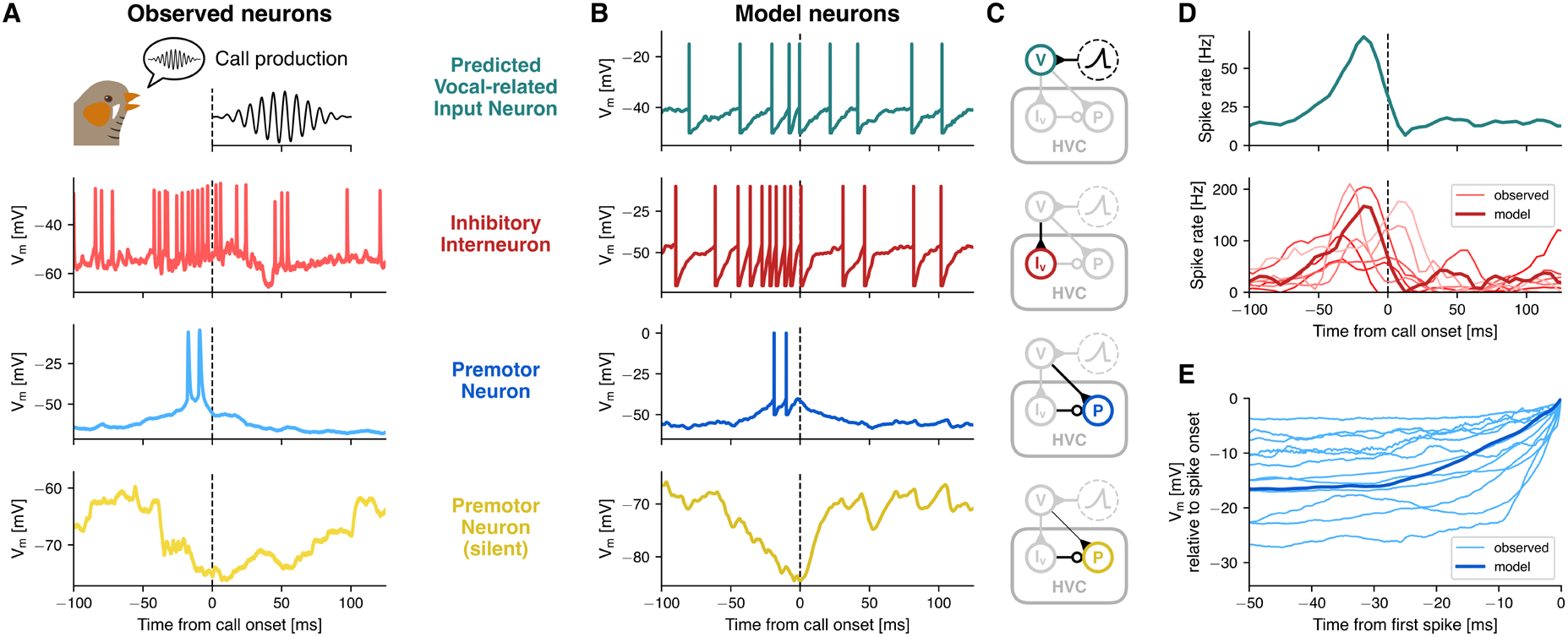
In silico call production-associated neural activity mirrors in vivo data. (**A**) Example membrane potential traces from intracellular recordings of an HVC inhibitory interneuron (red), a bursting HVC premotor neuron (blue) and a silent HVC premotor neuron (yellow) aligned to the onset of a call (dashed line) produced by the observed bird (data from Benichov & Vallentin, 2020). (**B**) Corresponding model traces of an interneuron (red), a bursting premotor neuron (blue) and a silent premotor neuron (yellow), as well as a neuron from a predicted population of upstream vocal-related input neurons (teal, top). (**C**) Circuit diagrams that show model connectivity and highlight the respective populations and their incoming connections. Neuron populations are represented as circles and synaptic connections between populations as lines ending either in excitatory synapses (triangles) or inhibitory synapses (circles). The predicted vocal-related population receives only a quadratically ramping input current that peaks and then returns to baseline prior to call onset (dashed circle). The silent premotor neuron receives the same input as the bursting premotor neuron, however excitatory weights from the vocal-related population are lower (8pA instead of 20pA). (**D**) Top: Spike rate of the predicted vocal-related population, aligned to call onset (dashed line). Bottom: Spike rate of seven intracellularly recorded interneurons that ramp up in activity prior to call onset, averaged across trials (light, thin lines), and average spike rate of the model interneuron population (dark, thick line). (**E**) Ramping subthreshold membrane potential of twelve intracellularly recorded HVC premotor neurons that burst around call onset (thin light blue lines) and the model premotor neuron (thick dark blue line). All traces were aligned to the time point and membrane potential of their first spike onset (set to zero). Recorded traces were averaged across trials and the model trace was averaged across 100 simulations, each with different randomized amplitude offsets in the input current onto the predicted vocal-related neurons.

In detail, the model consisted of an upstream population of 150 excitatory neurons (Mackevicius et al., 2020; Otchy et al., 2015; Danish et al., 2017), that projected onto both the premotor neuron and a population of 30 local inhibitory interneurons (Coleman & Mooney, 2004) (Figure 1B & C). Similar results were obtained with lower and higher numbers of neurons in those populations, as long as their ratio was around 5:1 (Figure S1). This predicted vocal-related population (“V”) was driven by a transient, ramping input current (Figure S2A). The resulting activity led to a transient increase in interneuron spiking (Figure 1D). The main features of the modelled interneuron activity captured the observed range of activity: simulated population activity peaked at 167 Hz (observed: 64.2 – 210.6 Hz), −17.5 ms relative to call onset (observed: −32.5 – 7.5 ms) and returned to baseline at 8.1 ms (observed: −15.0 – 52.4 ms). The vocal production-related input to the bursting premotor neuron also replicated the gradual increase in subthreshold membrane potential prior to the burst, which was observed in the intracellular recordings (Figure 1E). The silent premotor neuron was hyperpolarized through inhibitory input from the interneurons (observed mean hyperpolarization onset = −52 ± 14 ms). Additionally, it received excitatory input from the vocal-related population, whereby synaptic weights were lower compared to the bursting premotor neuron (Table 2). The longer duration of the hyperpolarization observed in the recorded neurons, compared to the model neuron, might be a result of receiving inhibition from multiple interneurons that reached peak activity at different time points (see Figure 1D).

### Feed-forward inhibition of premotor activity as a mediator of response timing

The described network model is biologically plausible, consisted of only a small number of components, and replicated observed call-related premotor and interneuron activity in the zebra finch HVC. The model is versatile and, considering what is known about the network components, there are several ways in which it can be interconnected. Here, we proposed three different model schemes and tested their relative ability to replicate previously observed changes in call production-related HVC activity and experimentally induced perturbations of the circuit.

In the first model, we assumed that inhibition does not play a functional role within HVC during call interactions (‘No Inhibition’ model, Figure 2A). Because the bursting premotor population in this network configuration was independent of any call-related inhibitory input from interneurons, it followed that its activity is unaffected by changes in the weights of inhibitory synapses (Figure 2B). Experimentally, however, we found that local disinhibition of premotor neurons through focal application of the GABA_A_ receptor antagonist, gabazine, resulted in stronger and earlier bursts relative to call onset (Benichov & Vallentin, 2020; Figure 2G). This discrepancy, together with evidence of the high connection probability between interneurons and premotor neurons in HVC (Mooney & Prather, 2005; Kosche et al., 2015; Kornfeld et al., 2017), suggested that the ‘No Inhibition’-model was insufficient as an explanation of call-related neural activity in HVC.

**Figure 2 –.**
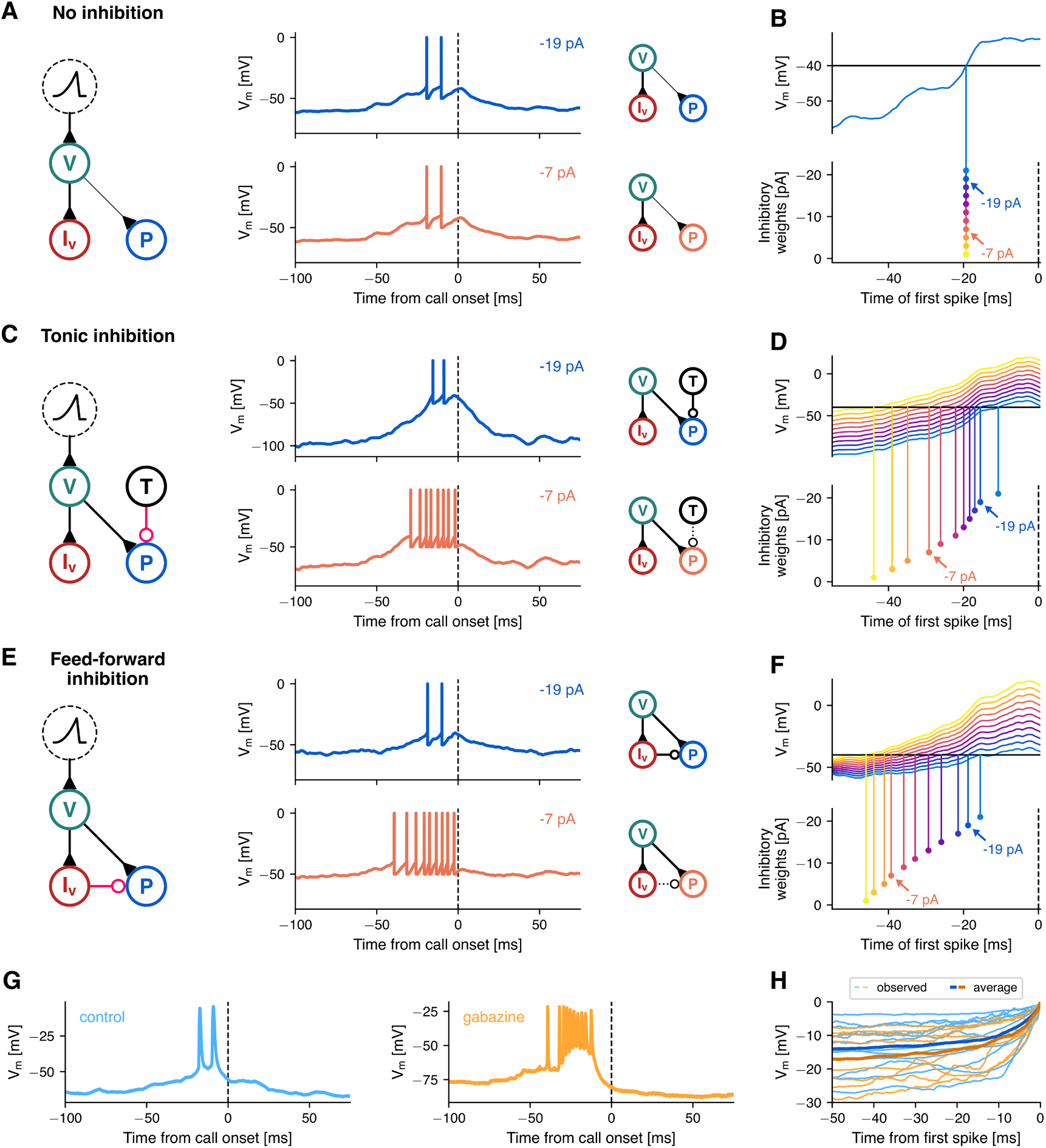
Feed-forward inhibition as a mediator for flexible response timing. Three alternative models consistent with the intracellular recordings in HVC (Figure 1), differing in connectivity (A, C, E). (**A**) In the ‘No Inhibition’-model, local interneurons (‘I_v_’, red) driven by vocal-related input neurons (‘V’, teal) silence a subpopulation of premotor neurons (not shown, see Figure 1B&C, bottom). Premotor neurons bursting prior to call onset (‘P’, blue) are triggered by input from the same vocal-related input population as interneurons. The traces show the membrane potential of the bursting premotor neuron when inhibitory weights were −21 pA (blue, top) and −7 pA (orange, bottom). (**B**) Membrane potential traces under a range of inhibitory weights, in the absence of a spiking mechanism. As bursting premotor neurons in this model do not receive inhibition, globally modifying inhibitory weights does not influence burst onset timing (i.e. spike threshold crossing). Arrows highlight the respective traces in (A). (**C-D**) In the ‘Tonic Inhibition’-model the bursting premotor neurons additionally receive inhibition from a population of tonically active interneurons (‘T’, black). Reducing inhibitory weights in this model pushes the membrane potential towards spike threshold, uniformly across time (i.e., both ramp and baseline potential). This leads to earlier premotor bursts, as well as a larger number of spikes per burst (bottom trace, −7 pA). (**E-F**) In the ‘Feed-Forward Inhibition’-model bursting premotor neurons receive feed-forward inhibition instead of tonic inhibition, partially balancing the excitatory ramping input. This allows inhibition to influence the steepness of the ramp and, to a lesser degree, impact the baseline membrane potential. As in (C-D), reduction of inhibitory weights leads to earlier and stronger premotor bursts. (**G**) Two example traces recorded from premotor neurons, one under control conditions (blue, same trace as in Figure 1) and one after microinfusions of GABA_A_ antagonist Gabazine in HVC (orange). Data from Benichov & Vallentin (2020). (**H**) Ramping subthreshold membrane potential of the observed HVC premotor neurons that burst around call onset in the control condition (light blue, n=12) and in after Gabazine microinfusions (light orange, n=8), as well as their respective averages (thick lines).

Next, we tested two models that incorporate inhibition, with a unidirectional local connectivity between interneuron and premotor neuron. As our focus was on the activity that resulted in a premotor burst, as well as the timing of these bursts, possible effects of premotor bursts through recurrent connectivity with interneurons were excluded. A direct inhibitory input to the bursting premotor neurons was added either in a tonic or phasic mode (the latter triggered by external inputs). Both temporal patterns of inhibition are biologically plausible and have been reported to maintain the excitatory/inhibitory balance of a network (Kosche et al., 2015; Vogels et al., 2011). In HVC, multiple types of interneurons have been characterized (Wild et al., 2005; Colquitt et al, 2021), exhibiting tonic firing patterns in vitro (Daou et al., 2013) and structured phasic activity during song production (Kosche et al., 2015).

The ‘Tonic Inhibition’-model included a population of consistently active interneurons synapsing onto the bursting premotor neuron (Figure 2C). In the ‘Feed-Forward Phasic Inhibition’-model, interneurons driven by the predicted vocal-related input neurons transiently affected bursting premotor activity (Figure 2E).

Both models simulated the activity patterns of premotor neurons and interneurons during call production. By simulating gabazine conditions through progressive reduction of the inhibitory weights on the premotor neuron synapses in 5pA steps, we asked how varying inhibitory weights influenced premotor burst onsets, strength, and subthreshold membrane potentials for each wiring scheme. In both models, premotor bursts occurred earlier and contained more action potentials, similar to the results obtained experimentally (Figure 2C–F; cf. Figure 2G).

The main difference between the Tonic Inhibition and Feed-Forward Phasic Inhibition models were apparent in the effects of inhibition on the membrane potential of premotor neurons preceding call-related firing. In the Tonic Inhibition model, inhibition acted equally across the entire peri-call interval. Therefore, reducing the weights effectively shifted the baseline membrane potential uniformly towards spike threshold. As a result, the ramping potential reached spiking threshold at successively earlier time points as the inhibitory synaptic weights are decreased (Figure 2D). On the other hand, in the Feed-Forward Phasic Inhibition model, the transient increase in interneuron firing counterbalanced the excitatory vocal-related drive during the pre-burst ramping more sparsely in time. In this case, when reducing inhibitory synaptic weights, we observed a more modest shift in the baseline potential as well as an increase in the steepness of the ramping subthreshold potential, resulting in an earlier threshold crossing and thus earlier and stronger premotor bursts (Figure 2F). The Tonic inhibition model, unlike the Feed-Forward Phasic Inhibition model, thus prediced a considerable increase in baseline membrane potential prior to premotor bursts caused by the reduction of inhibition, which was not observed during the experimental perturbation with Gabazine (Figure 2H & S3).

Taken together, these simulations demonstrate that a feed-forward connectivity between interneurons and premotor neurons was an effective way to capture the call-related activity data observed in experiments. To assess the model’s sensitivity to variations in parameter values, we ran simulations with a range of synaptic weights and population sizes for the excitatory and inhibitory inputs onto the premotor neuron. We tested the resulting premotor traces for consistency with two features observed in the electrophysiological recordings: a baseline membrane potential between 5 and 25 mV below spike threshold (Figure 1E) and the emission of 1 to 6 action potentials in the 50 ms preceding call production (Benichov & Vallentin, 2020). Those criteria were fulfilled in a relatively broad range of synaptic weight combinations (Figure S4) and population sizes (Figure S1). Reduction of excitatory weights in this Feed-Forward Phasic Inhibition model could cancel and ultimately reverse the effect of the ramping input, leading to a hyperpolarization of the premotor neuron (Figure 1B & S4). Reducing the inhibitory weights, on the other hand, resulted in both stronger and earlier premotor bursts, suggesting a role of HVC interneurons in call timing control which could be confirmed in future experiments.

### Heterogeneity in HVC and its inputs’ activity profiles suggests an additional source of call-related fast inhibition

The Feed-Forward Phasic Inhibition model recapitulated the neural activity in HVC that generates motor output, i.e. precise premotor burst associated with call production. To enable alternating vocal turn taking, the bird does not only have to produce a call but also listen to its vocal partner. Therefore, we explored the neural dynamics during auditory perception and tested the robustness of the Feed-Forward Phasic Inhibition model in capturing sensory-related neuronal responses. To this end, we first looked for auditory-evoked activity in HVC by performing extracellular recordings in four awake, head-fixed (and as a consequence, vocally unresponsive) zebra finches (n = 225 neurons) while presenting a set of call playbacks. Under these conditions, responses observed in HVC are less likely to be confounded by activity that is directly related to vocal production.

Sparse bursting activity of premotor HVC_(RA)_ projection neurons in adult zebra finches only occurs during song or call production (Hahnloser et al., 2002). In addition, HVC projection neurons have been shown to be unresponsive to song playback in awake adult zebra finches (Vallentin & Long, 2015) whereas interneurons increase their activity in response to the tutor song presentation (Vallentin et al., 2016). We therefore hypothesized that our neural recordings in response to call playbacks were oversampling the activity of HVC interneurons. To test this, we classified neurons as putative interneurons or projection neurons based on spike waveform features (Figure 3A). This analysis showed an overrepresentation of neurons (171/225 units) with fast (trough to peak = 0.1955 ± 0.0422 ms) and narrow waveforms (FMHW = 0.4005 ± 0.0749 ms), characteristic of interneuron populations (McCormick et al., 1985) (Fig 3A; marked in black after k-means clustering).

**Figure 3 –.**
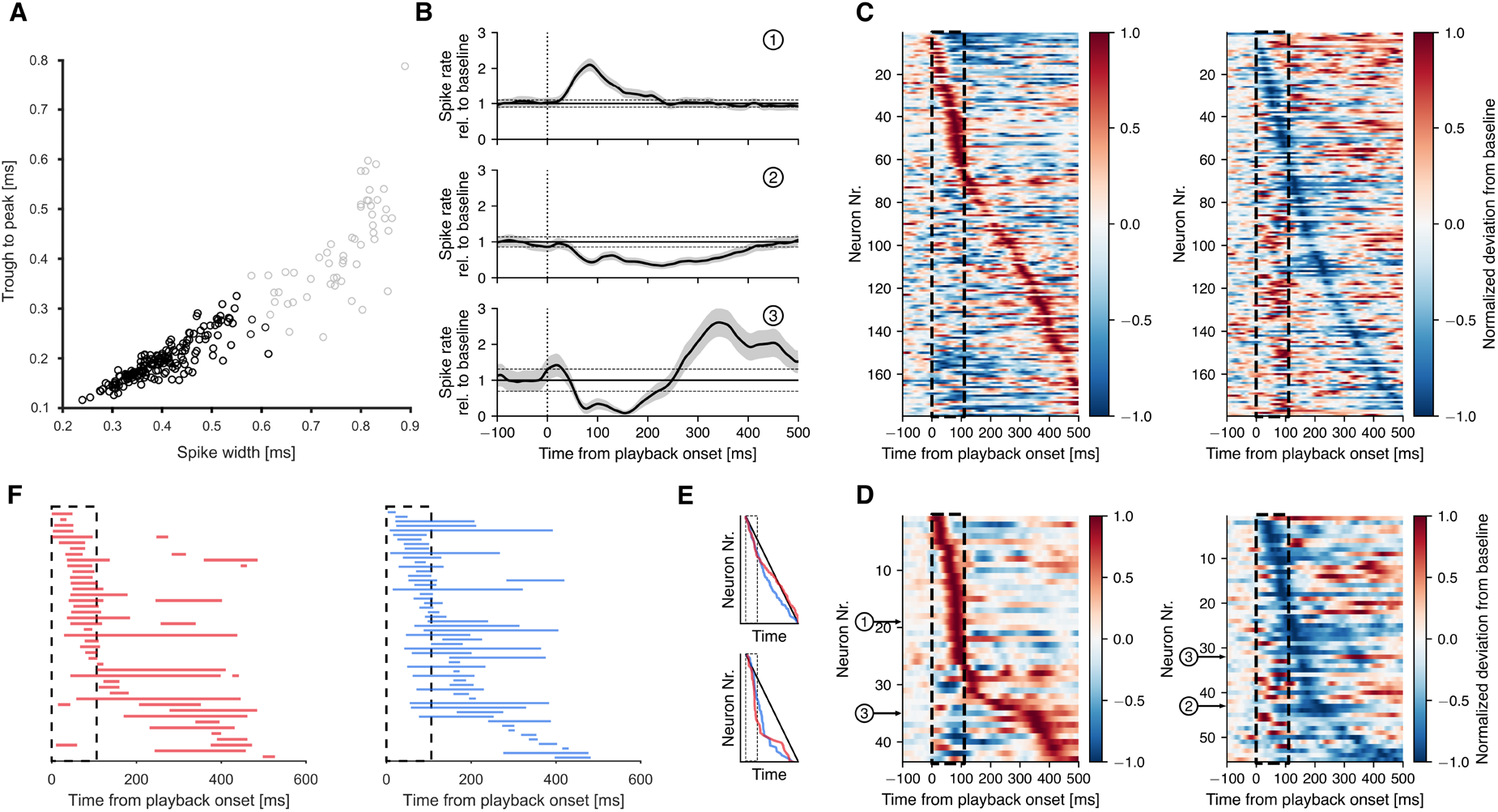
Responses in HVC to call playback stimuli. (**A**) Distribution of spike waveforms in feature space. Black dots are putative interneurons, determined by k-means clustering. (**B**) Average spike rate of three HVC neurons in response to call playbacks, normalized to baseline activity. Example of excitatory (top), inhibitory (middle) and mixed response (bottom). Gray patches mark average ± SEM. Horizontal lines mark baseline activity ± 2 standard deviations. Significant responses were defined as periods in which average rate - SEM exceeds baseline activity + 2 standard deviations and vice versa. (**C**) Average spike rates from all cells recorded for a minimum of 20 trials, normalized to baseline (0) and absolute maximum deviation from baseline (1 or −1), aligned to call playback onset and sorted by time of maximum positive (left) or negative deviation (right) after playback onset. Dashed black lines mark time of call playback. (**D**) Subsets of neurons that show significant excitatory (left) and/or inhibitory responses (right) after playback onset, sorted by peak of positive or negative deviation, respectively. (**E**) Time of the sorted positive (red) and negative peaks (blue) as seen in (C) (top) and (D) (bottom), compared to the values expected if peaks were distributed uniformly in time (black diagonal line). (**F**) Onsets and offsets of significant excitatory (left) and inhibitory (right) responses per neuron, sorted as in (D).

For further analysis, we took only neurons into account which were recorded during the presentation of at least 20 playbacks (179/225 units; Figure 3C). We classified neurons to be call-responsive when their average activity ± SEM after call playback onset crossed a threshold of two standard deviations above/below baseline firing rate (79/179 units; Figure 3B & D). We were able to distinguish three general neural response patterns among the call-responsive neurons: increases in firing activity after call-playback onset, suppression of firing in response to the playback stimulus complex mixed responses with excitatory and inhibitory phases (Figure 3B). We explored this response heterogeneity in call-responsive cells in further detail by calculating the response onset (126.94 ± 123.38 ms) and response duration (104.69 ± 98.02 ms). We found that cells can exhibit increased activity during call-playbacks with relatively fast response times or delayed increased activity after the offset of playbacks (Figure 3B). Suppressive responses occurred rapidly after playback onsets or with delayed onsets after playback offsets. This suggests that calls of a vocal partner can provide fast or delayed excitatory inputs onto HVC which can drive increases as well as decreases in HVC interneuron activity. When sorting the neurons that showed an excitatory response by the timing of their maximal firing rate exhibited during and after call playback we observed a strong overrepresentation of excitatory responses during playback and an underrepresentation in the 200 ms following playback offset (Figure 3D, left; 3E bottom). In contrast, when sorting the neurons that showed an inhibitory response by the timing of the peak of their minimum firing rate, we saw an overrepresentation of inhibitory responses from playback onset until 100 ms after playback offset (Figure 3D, right; 3E bottom).

Since HVC is primarily a premotor nucleus generating the timing of vocal output, we wondered whether the call playback-evoked excitatory response might have the potential to directly trigger vocal production. To better understand the auditory signals that contribute to call-related activity in HVC, we investigated its main source of higher auditory input - the Nucleus Interfacialis (NIf) (Lewandowski et al., 2013). We presented call playbacks to the awake bird while performing intracellular recordings of NIf neurons (n = 6 identified HVC projection neurons, 1 non-identified). These NIf_(HVC)_ neurons displayed call-related activity represented by either a suppression (−2.6 ± 3 Hz delta from baseline (silent period 300 ms prior to playback) or increase in firing rate (5.6 ± 6.6 Hz delta from baseline) in response to call playbacks (Figure 4A). Call-related activity was fast (onset time 35 ms) and the HVC neurons are a plausible recipient of this information (Coleman & Mooney, 2004). In line with similar response patterns to long distance call playbacks previously observed in unidentified NIf cells (Lewandowski, 2011) we hypothesized that NIf activity contributes to call-related activity changes observed in HVC.

**Figure 4 –.**
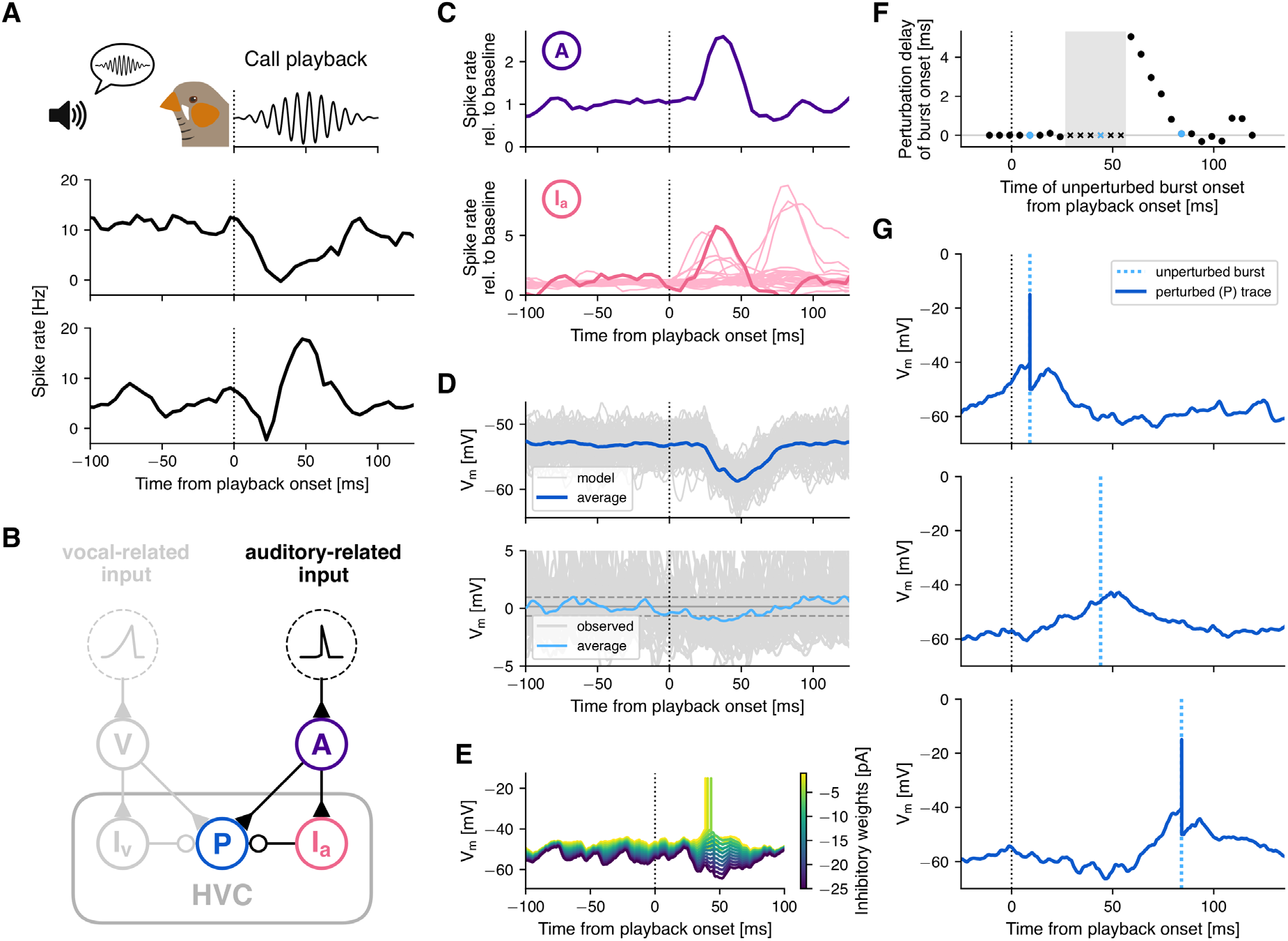
Playback-evoked inhibition can suppress premotor bursts associated with an imminent call. (**A**) (**B**) Circuit diagram of the feed-forward inhibition model, expanded with an auditory-related input population (‘A’, purple) and a second inhibitory population (‘I_a_’, pink) providing excitatory and feed-forward inhibitory input to the premotor neuron (‘P’, blue), respectively. (**C**) Population activity of the model auditory-related input population (top), which receives a short, ramping input current (see Figure S2) after call playback onset (dotted line). This triggers a peak of activity in the interneuron population (bottom), that is consistent with putative interneurons recorded extracellularly in HVC (neurons that significantly increased their activity within 100 ms after playback onset; light pink). (**D**) The bursting premotor neuron (‘P’, blue in (B)) at rest (i.e. while not receiving vocal-related input from V & I_v_) is transiently hyperpolarized. Model traces from 100 simulations with different randomized input currents (gray) and the average (blue). Below: Intracellular recordings of an example premotor neuron aligned to playback onset (dotted line), which is significantly hyperpolarized following playback onset. Horizontal lines show mean baseline potential ± 2 standard deviations (baseline: −100–0 ms). (**E**) Reduction of inhibitory weights onto the model premotor neuron reverses its playback-induced hyperpolarization (see D), ultimately eliciting a spike. (**F**) Simulation of the interaction between pre-call premotor activity (ramp and burst) and playback-evoked inhibitory suppression at different relative time points. Premotor bursts can be suppressed (marked by crosses) or delayed (y-axis) by playback-induced inhibition when the premotor burst occurs at different time points (x-axis) relative to the playback onset (dotted line). The gray rectangle marks a time window during which premotor bursts are suppressed. (**G**) Three example traces from the premotor neuron, bursting at different times relative to call playback. Top: Burst occurs before peak in inhibition and is therefore not perturbed relative to the burst onset without inhibitory suppression (blue dotted line). Middle: Burst is suppressed as pre-burst ramp occurs during inhibitory suppression, hindering the membrane potential to reach spike threshold. Bottom: Ramp is modified by inhibition, but potential still reaches threshold after a minimal delay.

Given that birds responded vocally with an average delay of ~200 ms to call playbacks, the function of the fast call-induced input from NIf we observed cannot fully explain the direct triggering of call responses. Instead, this activity has the potential to drive inhibitory interneurons in HVC which do not directly play a role in eliciting vocal output. Inhibitory interneurons in HVC synapse locally onto premotor neurons and this additional fast source of local inhibition in HVC could serve to suppress premotor activity of an imminent call. This mechanism could thereby reduce the likelihood of call overlap.

### Inhibitory suppression of premotor activity can reduce call overlap

Next, we investigated the interaction between call-related premotor drive and auditory-evoked inhibition. To do so, we extended the Feed-Forward Phasic Inhibition model (Figure 1 & 2E) with a second set of upstream excitatory (‘auditory’) and local interneuron populations. These were connected through the same circuit motif of excitation and feed-forward inhibition (Figure 4B). Based on the synaptic delay of auditory input (Margoliash, 1983) and the observed activity profile within NIf (Figure 4), the auditory population received a shorter ramping input current that peaks at 35 ms after playback onset with a short quadratic upstroke and linear downstroke (Figure S2B). In the interneuron population, this led to a transient peak in activity that matched the observed activity of a subset of the putative interneurons (Figure 4C). In contrast to the original model (Figure 2E), the balance of synaptic weights in the auditory model is biased towards inhibition, so that the premotor neuron in the absence of ramping input from the original vocal-related population was transiently hyperpolarized after call playback. We observed a similarly timed, albeit weaker hyperpolarization when we aligned intracellular recordings of premotor neurons to playback onset (Figure 4D). Other premotor neurons were slightly depolarized in the same timeframe (Figure S5).

To simulate call production at different time points relative to the playback, we varied the time difference between the input currents to the vocal-related and the auditory population while observing the delay or the suppression of bursts caused by auditory-evoked inhibition. Between 25 and 55 ms after playback onset, bursts were suppressed, as playback-evoked inhibition prevented the neuron from reaching spike threshold (Figure 4F & G). Bursts which occurred earlier were unaffected, while later bursts were delayed by up to 5 ms, due to a perturbation of the pre-burst ramp in subthreshold potential.

After we determined the time window following playback onset, during which premotor bursts were suppressed (‘suppression window’: 25 – 55 ms), we next estimated a time window prior to the onset of call production, during which the suppression of premotor bursts can potentially cancel the imminent call (‘estimated window of susceptibility’). Average burst onset time of the observed premotor cells varied between −45.0 and +33.4 ms relative to onset of call production (Figure 5B). We set the start of the estimated window of susceptibility to 60.95 ms before call onset (mean - standard deviation of the earliest average burst onset, Figure 5B). The end of the window was defined as 10 ms prior to call onset, as we assumed that after this time point any further changes at the level of HVC cannot influence call production, while the call was initiated further downstream.

**Figure 5 –.**
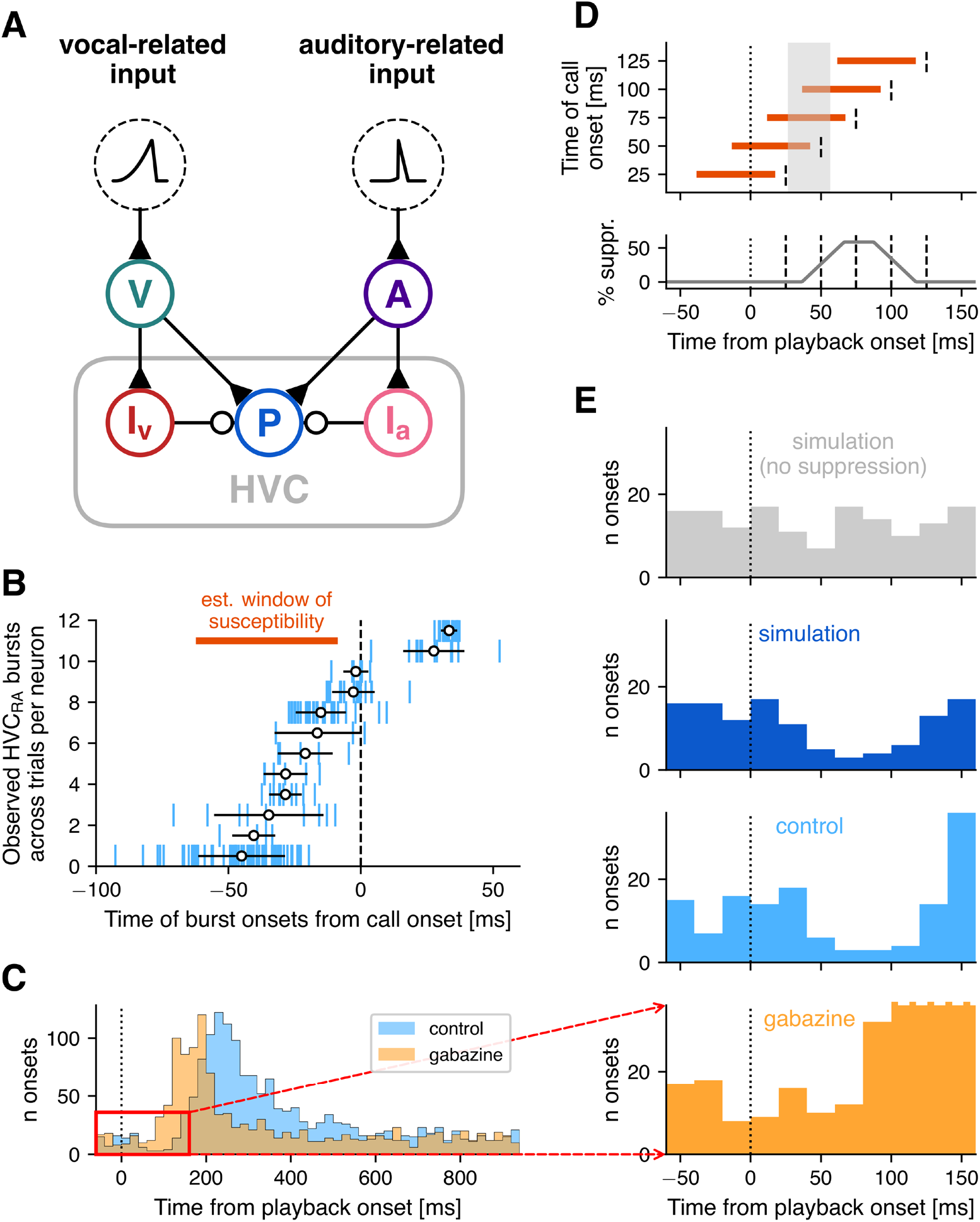
Playback-evoked inhibitory suppression of premotor activity can reduce call overlap. (**A**) Circuit diagram of the full model. (**B**) Timing of all burst onsets during multiple trials relative to onset of call production (x-axis) for all 12 observed HVC premotor neurons in the control condition (y-axis). The orange bar marks the estimated time window during which premotor neurons triggering call production are susceptible to inhibitory suppression. (**C**) Full histograms of call onsets during control (blue) and Gabazine conditions (orange) across the one second inter-playback interval. The red rectangle highlights the section depicted in (E). (**D**) Five example call-onsets (dashed lines) with their associated windows of susceptibility (orange bars). Below is a function of the percentage of overlap of the window of susceptibility with the suppression window (gray rectangle, see Figure 4F), i.e. the percentage of premotor suppression (y-axis) against the onset time of a hypothetically produced call. Example call onsets from above are marked by dashed lines. (**E**) We simulated a random uniform distribution of call onsets (gray) and removed calls with a likelihood proportional to the suppression function in (D). The resulting call onset distribution (blue) matches that of the behavioral experiments in the control condition (light blue), both of which show a dip in call likelihood between around 50 and 110 ms after playback onset. Consistent with our model, Gabazine microinfusions in HVC abolished that dip (orange). Instead, response likelihood begins to sharply increase around 80 ms after playback onset. Histogram bars after 100 ms are cut off in the Gabazine panel. Data in (B, C & E) from Benichov & Vallentin (2020).

Relating the suppression window with the estimated window of susceptibility allowed us to predict the behavioural outcome of the proposed suppression mechanism, given two assumptions regarding the temporal distribution and function of the pre-call premotor drive. First, we assumed that bursts were distributed nearly uniformly across time before call onset. Such a distribution has long been hypothesized for premotor neurons during song production (e.g., Hahnloser et al., 2002; Fee et al., 2004; Long et al., 2010) and more recently received support from electrophysiological recording and imaging of large populations of HVC projection neurons (Lynch et al., 2016; Picardo et al., 2016). Despite a smaller dataset, intracellular recordings during call production suggested a similar distribution for pre-call activity (Figure 5B).

Second, we assumed that the triggering of a timed call response depends on the number of premotor spikes. Call-like vocalizations can be elicited by electrical stimulation of down-stream nucleus DM, however only above a certain stimulation threshold (Vicario & Simpson, 1995, Ashmore et al., 2008; Fukushima & Aoki, 2000). It is likely that excitatory input to DM from HVC (via RA) is sufficiently strong to elicit a call response. Suppression of a significant number of premotor bursts through auditory-evoked inhibition would thus reduce the likelihood of a call response. For our model we decided to assume a linear relationship between the magnitude of premotor activity and the probability of call initiation.

Given these two assumptions, the amount of overlap between the suppression window and the estimated window of susceptibility predicted the likelihood of a call being triggered. We determined the overlap as a function of call onset timing relative to playback onset (‘suppression function’, Figure 5D). If a bird were to call at random times in the absence of playback, we would expect playback-evoked inhibition to result in a dip in the call onset distribution shortly after the onset of playback. We first generated uniformly distributed random call onset times (Figure 5E, grey). To simulate playback-evoked call suppression, we then removed each call with a probability proportional to the suppression function (Figure 5E, blue, see Methods).

Through this process we effectively simulated the behavioral output (i.e., call response time distribution) predicted by the modeling of fast and transient auditory-evoked inhibitory suppression of premotor activity. According to the prediction, call likelihood decreases between 50 and 110 ms post playback. Inhibitory suppression in the model had the potential to suppress call production shortly after an incoming auditory cue, and could thereby partially reduce the overlap of calls between two vocally interacting birds. Complete overlap of calls (i.e., two birds initiating a call within 50 ms of each other) was not affected, as in this case the initiation of each call would occur before auditory information about the partner’s call affects activity in HVC.

For a comparison to observed behavioral data (Benichov & Vallentin, 2020), we pooled the call onset times of all birds responding to a regularly timed call playback (one call per second) in either a control condition or after gabazine application (Figure 5C). The onset of call suppression in the predicted call onset distribution matches that of the control condition (Figure 5E). At around 150 ms after playback onset, the observed call responses sharply increased above the pre-suppression baseline. At this point the predicted distribution deviated from the recorded distribution, as increased call likelihood in response to the playback was not factored into the model. In the gabazine condition, no reduction of call responses following playback could be observed (Figure 5C & E). This outcome was expected, as a reduction of inhibitory efficacy in HVC reduces or even eliminates the effect of the proposed suppression mechanism. Instead, response likelihood increased above baseline between 80 and 90 ms after playback onset, i.e. already before playback offset.

Taken together, these results indicated that inhibition within HVC regulated the behavioral output on two time scales: On a short time scale, an auditory-evoked increase in inhibition led to a suppression of vocal motor output while the social partner was producing a vocalization and, thus, a call was being withheld and vocal overlap prevented. On a longer time scale, inhibition was related to the premotor preparation and controlling the precise timing of a vocalization.

### Inverting excitatory/inhibitory balance leads to auditory triggering of calls instead of suppression

The observed fast call responses cannot be explained by the vocal-related input alone. When reducing feed-forward inhibitory weights in the model, premotor bursts occur, at most, 50 ms earlier (Figure 2F). While this timescale could partially account for the reductions in call response latency observed in the gabazine experiments, it did not fully explain the observation of the largest time differences in the case of the fastest responses (the most extreme bird reduced its response latency by 200 ms after Gabazine application; Benichov & Vallentin, 2020).

Having validated this model for auditory-evoked suppression of call production, we thus wanted to test whether the fastest responses during gabazine treatment could be directly triggered by auditory-related input. Therefore, we gradually decreased the synaptic weights of the auditory-driven interneuron population onto the premotor neuron, mimicking the effects of gabazine application. As the inhibitory weights decreased, the excitatory drive of the auditory population increasingly dominated the synaptic input, leading to a transient depolarization in the premotor neuron (Figure 4E). With reduced inhibitory weights (< 6 mV), the auditory input elicited a spike.

## Discussion

We developed a network model of zebra finch HVC that illustrates how cortical control over innate vocalizations (calls) can facilitate vocal turn-taking. In the proposed model, HVC integrates auditory and premotor information and gates the production of call responses by sending excitatory input to downstream vocal-motor nuclei at appropriate times.

The model accounts for the observation that the restriction of inhibitory influence in HVC leads to birds responding significantly faster to the calls of a vocal partner (Benichov & Vallentin, 2020). This reduction in response latency can be brought about by the shift in balance of excitatory and inhibitory input onto model premotor neurons in two (non-exclusionary) conditions: fast and slow responding. First, the dominance of excitation during the integration of vocal-related input causes premotor neurons to reach spike threshold earlier, predicting a reduction in response latency on the order of 50 ms. Second, if the fast auditory-evoked neural response (< 50 ms) to call playback leads to a strong enough depolarization, it can lead to premotor spiking activity even before the arrival of production-related input. Whether this is the case in vivo, and whether this activity would suffice to trigger a call response remains to be investigated.

One prerequisite for the replication of our model of the in vivo recorded activity of HVC neurons during calling is an excitatory “vocal-related” input to HVC occurring at the onset of call production-related changes in activity. This raises the question: What is the source of excitation that would drive an increase in interneuron activity and causes premotor neurons to burst?

For calls that are produced in response to the heard calls of conspecifics, afferent auditory-related input onto HVC would be one likely source. It is known that premotor nucleus HVC receives excitatory input from multiple areas: the thalamic nucleus UVA sends both vocal- and auditory-related information to HVC (Hahnloser et al., 2008; Danish et al., 2017; Akutagawa & Konishi, 2005). Sensorimotor nucleus NIf (Nucleus interfacialis) provides the largest source of auditory information onto HVC premotor neurons and interneurons (Coleman & Mooney, 2004; Rosen & Mooney, 2006; reviewed in Lewandowski et al., 2013), and there is evidence of direct auditory input from other regions of the auditory forebrain as well (Shaevitz & Theunissen, 2007).

Although there is some evidence for direct input from auditory forebrain areas Field L and the lateral caudal mesopallium (CM; Shaevitz & Theunissen, 2007), NIf appears to be a likely candidate area for several reasons. NIf projects directly onto HVC and provides its strongest source of auditory information (Lewandowski et al., 2013; Janata & Margoliash, 1999; Cardin & Schmidt, 2004). The time course of activity of the predicted vocal-related input population in relation to the onset of calls (Figure 1D) closely matches that of neurons previously recorded in NIf during call production (Lewandowski, 2011). The timing of call-related NIf activity relative to call-related activity in HVC is consistent with monosynaptic inputs.

While call-related increase in interneuron activity necessitates an excitatory drive, premotor bursts could hypothetically be a result of post-inhibitory rebound depolarization. However, this phenomenon appears to be absent in most premotor neurons in adult zebra finches (Daou et al., 2013; Ross et al., 2017, 2019), reducing the likelihood that premotor bursts were triggered solely by the offset of inhibition, without any excitatory input. Another excitatory neuron type in HVC that projects to “Area X” of the basal ganglia does exhibit rebound spiking. These cells sparsely synapse onto premotor neurons (Mooney & Prather, 2005) and could thereby theoretically induce premotor bursts in a scenario in which external excitation only drives interneurons (Ross et al., 2017). interneurons, however, do not return to their baseline firing rate until after call onset 20.9 ± 19.9 ms which is after the average burst onset of premotor cells (−14.4 ± 23.8 ms). Thus, the relative timing of premotor and interneuron activity and the sparse connectivity profile between HVC-X neurons and premotor neurons does not support rebound spiking induced excitation as a mechanism for premotor drive.

It is important to note that we modelled a single hypothetical bursting premotor neuron, which we assume to be representative for the entirety of premotor neurons. The recorded activity among the different premotor and interneurons was qualitatively similar: sparse bursts and a transient increase in firing rate, respectively (Benichov & Vallentin, 2020). Each individual neuron exhibited a relatively stereotyped time course across trials, with respect to call onset. Across neurons, however, the timing differed for both premotor and interneurons (Benichov & Vallentin, 2020; Figure 1D & 5B). Similar variability in the timing of vocal-related input neurons could account for these observations. Subsets of these neurons that ramp up in activity at different time points could thus drive different subsets of HVC premotor and interneurons that become active at different time points relative to call onset.

In conclusion, the model we propose allowed us to examine social coordination from the perspective of a relatively simple sensorimotor circuit and has highlighted several potentially important mechanisms. Specifically, vocalization-related premotor inhibitory strength can achieve temporal fine-tuning of vocal responses and auditory-evoked inhibition can temporally suppress premotor drive, thereby reducing simultaneous calling, e.g. ‘jamming’. The role of inhibition, in both of these regulatory processes, is more extensive than previously thought and suggests that further investigation of inhibitory cell types and connectivity are required within the songbird vocal-motor pathway and other sensorimotor circuits more broadly. The underlying feed-forward wiring scheme of excitatory and inhibitory neurons can be found across brain areas and species. Applying this model to the study of vocal turn-taking in other experimentally tractable model systems, including singing mice (Okobi et al., 2019) and marmosets (Takahashi et al., 2013, 2016; Dohmen & Hage, 2019), would determine if these mechanisms are general inhibitory principles of interactive vocal control. Our model therefore provides a versatile framework for testing predictions about vocal turn-taking behaviors observed across a variety of times scales and species.

## Materials and methods

### Animals

All animal care and experimental procedures were performed with the ethical approval of the Max Planck Institute for Ornithology and the Regierung von Oberbayern (ROB-55.2-2532.Vet_02-18-182). For extracellular recordings, we used 4 adult male zebra finches (> 90 days post hatching) that were acquired from the breeding facility at the Max Planck Institute for Ornithology. –Throughout the experiments, the birds were maintained in a temperature and humidity controlled environment with a 14/10 hour light/dark schedule and ad-libitum food and water.

### Surgery

Zebra finches were anesthetized with isoflurane (1–3% in oxygen). The centers of RA and HVC were located based on stereotactic coordinates and two small craniotomies were performed at the targets. Intracellular microdrive implantation and pharmacological perturbations were previously described in Benichov & Vallentin 2020. In all cases, a chlorided silver ground wire (0.001”, California Fine Wires) was implanted above the cerebellum. For antidromic identification of HVC-RA projecting premotor neurons, a bipolar stimulating electrode was implanted into the downstream nucleus RA. A custom-made stainless steel head plate was affixed to the skull using dental acrylic (Paladur, Kulzer International). The craniotomies were protected until experiments were conducted using a silicone elastomer (Kwik-Cast; WPI). Animals were returned to their home cage with a companion bird for at least 24 hours post-surgery and were monitored to ensure full recovery before experiments commenced.

### Playbacks

For measuring neural responses to calls, we presented call playbacks at 65 dB through a speaker placed in front of the head-fixed birds. The presented stimuli were recordings of an average male “stack” call, presented using a custom-made labview interface in blocks of 10 at a rate of 1 call per second (Benichov et al., 2016), with 3 seconds of silence between each block. A 10 ms 15 kHz pulse (beyond zebra finch auditory range) was simultaneously played at the onset of each stimulus to ensure subsequent uniform alignment of playbacks with the neural data.

### Electrophysiological recordings

Awake-behaving intracellular microdrive recordings were performed as previously described in detail (Benichov & Vallentin, 2020). Extracellular recordings during call playback were performed in head-fixed awake birds held in a soft foam restraint. A 16-channel silicon probe (NeuroNexus) was lowered into HVC (between 300-700 μm from the dorsal surface) using a micromanipulator (Sutter Instruments). Neural activity was digitized at a sampling rate of 30 kHz on an Intan RHD2132 headstage and acquired with an RHD Recording Controller (Intan Technologies). A TTL pulse was triggered by the 15 kHz tone at the onset of each playback presentation using an Arduino Uno, and delivered to the RHD Recording Controller for acquisition alongside the neural data.

### Data Analysis

We used Plexon Offline Sorter for spike detection and clustering and MATLAB R2020a and Python 3.7 for data analysis. For the analysis of the extracellular recordings (Figure 3) only neurons were regarded that had a minimum of 20 trials (i.e. playbacks).

Spike rate time series in Figures 1D and 4A & C were calculated with a bin size of 5 ms and smoothed using a Savitzky-Golay filter with window length 9 and polynomial of order 3 (. Spike rate time series in Figure 3 were calculated with a bin size of 11.1 ms, linearly interpolated to a 1 ms resolution and then smoothed using a Savitzky-Golay filter with window length 99 and polynomial of order 2.

Significant responses of the extracellularly recoded HVC neurons (Figure 3) were determined as follows: Responses were defined as periods in which the average spike rate ± SEM after call playback onset crossed a threshold of two standard deviations above/below baseline firing rate remained above/below this threshold for at least 15 ms. If two positive or negative response onsets followed each other within a time interval of 200 ms, the two responses were merged and counted as one response starting at the onset of the first and ending at the offset of the second response. If the gap between two positive or negative reponses was shorter than 120 ms, then this time interval was extended to 350 ms.

One of the eight intracellularly recorded interneurons was omitted from the analysis (Figure 1D and corresponding values), due to the low number of trials (n=3). Its activity peaked at 59.2 Hz, 2.5 ms after call onset and returned to baseline at 71.6 ms.

### Neuron model

To simulate the membrane potential dynamics of neurons in the zebra finch song system, we used a leaky integrate-and-fire neuron model with current-based synapses. The voltage dynamics of the membrane are described by the equation

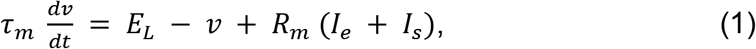

where *v* is the membrane potential, *E_L_* the leak potential (or resting potential), *R_m_* the membrane resistance, and *τ_m_* the membrane time constant. When the membrane potential of a neuron reaches its threshold *v_thresh_*, it is instantaneously set to its reset potential *v_reset_* and a spike is emitted. *I_e_* and *I_s_* are the electrode current and synaptic current, respectively. *I_e_* is used to inject either a time-varying current into the predicted ustream populations (Figure S2) or a small constant current representing unspecific background excitation. Synaptic currents are determined by:

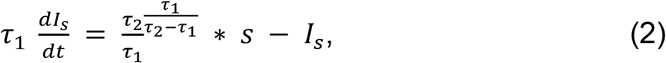

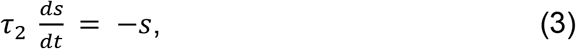

where *τ*_1_ is the decay time constant (*τ_decay_*) and *τ*_2_ the rise time constant (*τ_rise_*) of the bi-exponential synaptic current (Figure S6). Each time a presynaptic neuron spikes, the corresponding synaptic weight is added to *s* in the postsynaptic neuron.

Parameters for excitatory and inhibitory model neurons and synapses were fit to data from electrophysiological studies of zebra finch HVC_(RA)_ premotor neurons and HVC interneurons, respectively (Mooney & Prather, 2005; Kosche et al., 2015; Hamaguchi et al., 2016), and are given in Table 1. As such studies are sparser for nuclei upstream of HVC, and as we don’t know the exact source of the excitatory input we propose, we chose to use the same parameters that were fit to HVC premotor neurons for the predicted upstream populations (“excitatory” in Table 1).

### Input currents

Neurons in the predicted upstream populations (vocal- and auditory-related input) are driven by a time-varying input current that is aligned to the onset of call production or playback onset, respectively. For the vocal-related input neurons, these currents are characterized by a constant baseline current, followed by a quadratic upstroke from 80 to 10 ms prior to call onset and a linear return to baseline from 10 to 0 ms prior to call onset (Figure S2A). Input to the auditory-related input neurons ramps up between 10 and 35ms after playback onset and returns to baseline between 35 and 60 ms (Figure S2B). For the vocal-related input neurons, baseline current is 170 pA and input peaks at 220 pA (auditory-related: 168 pA and 180 pA, respectively).

To induce variability between neurons in their spiking pattern, at each time-step a timevarying offset is multiplied with the current value at that timestep and added to the input current. This time-varying offset changes every millisecond, where a new pseudorandom value is drawn from a normal distribution with mean 0 pA and variance 200 pA (Figure S2). Additionally, each current segment (baseline, upstroke, downstroke, baseline) for each neuron is offset by a pseudorandom value drawn from a normal distribution with mean 0 pA and variance 10 pA. The remaining neural populations (inhibitory and premotor neurons) receive a constant input current of 30 pA that represents unspecific background excitation.

### Network connectivity

Model neurons between the different populations are connected randomly in an all-to-all manner, with connection probabilities given in Table 2. There are no recurrent connections between neurons within populations.

### Simulation

Model simulations were carried out in Python 3.7 using Brian 2 version 2.2.2.1 (Stimberg et al., 2019). Equations (1–3) were integrated analytically (using Brian’s ‘exact’ method), with a constant time step of 0.02 ms.

Except for the premotor neuron, all model neurons were initialized with different membrane potentials between *E_L_* and *v_thresh_*, drawn pseudorandomly from a uniform distribution.

For visualization purposes, artificial spikes were added to the model voltage traces in Figures 1, 2, 4 and 5 as vertical lines above spike threshold at the time points of each spike.

## Supplements

**Figure S1 –.**
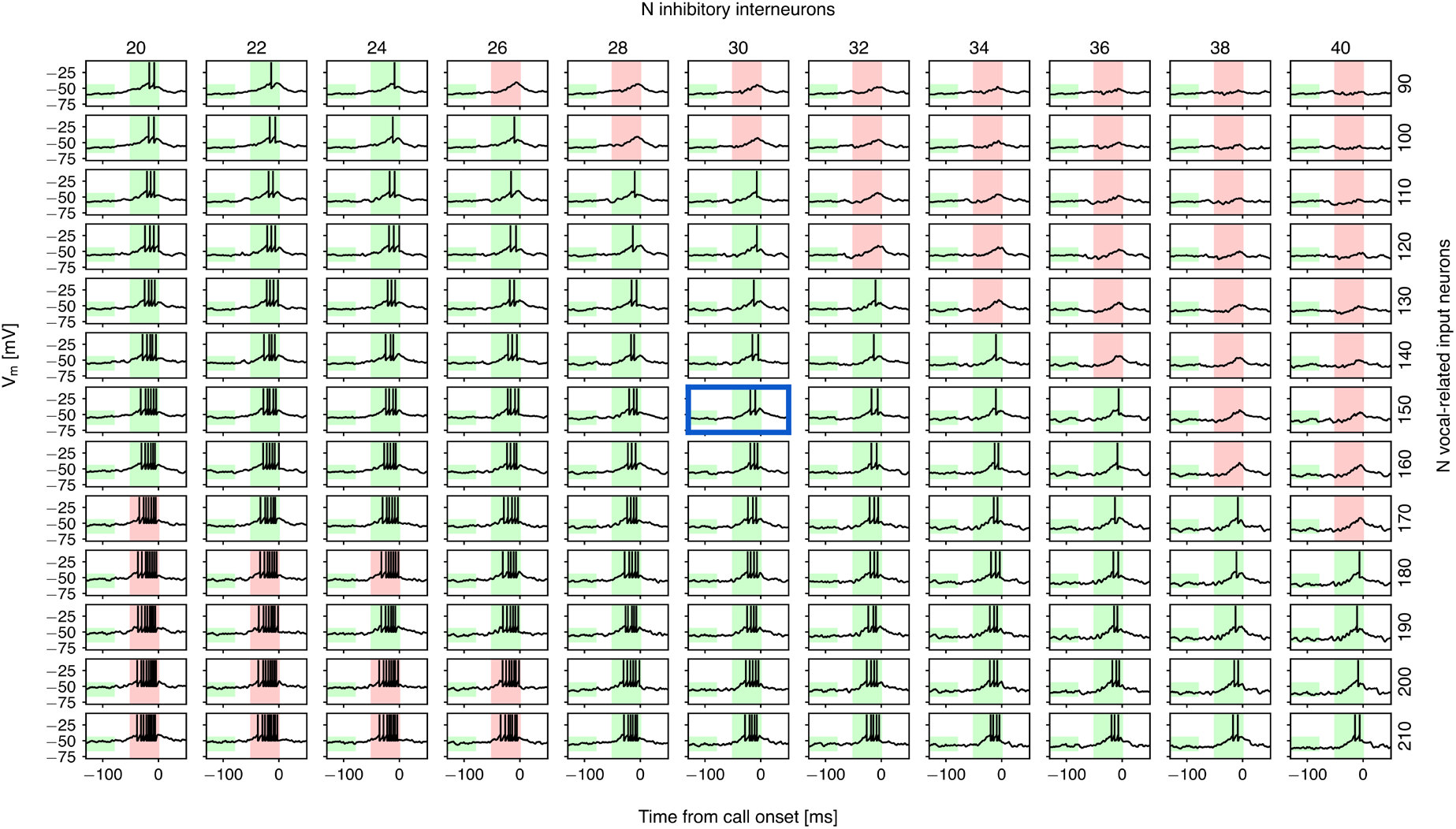
Sensitivity analysis for the Feed-Forward Phasic Inhibition model: population sizes. Membrane potential traces of the model premotor neuron in dependence on the number of inhibitory interneurons (left to right) and vocal-related input neurons (top to bottom). The colored rectangles show whether two criteria are fulfilled (green) or violated (red) in the different simulations: First (left rectangle in each panel), that the baseline membrane potential (average potential between 130 and 80 ms prior to call onset) is in the range of the recorded premotor neurons. This range is between 5 and 25 mV below spike threshold (see e.g. Figure 1E), which corresponds to − 65 – −45 mV in the model. Second (right rectangle in each panel), that the neuron produces between one and six spikes during the 50 ms prior to call onset, as was observed in the recorded premotor neurons. The blue frame marks the parameter combination used in the simulations (30 interneurons, 150 vocal-related input neurons).

**Figure S2 –.**
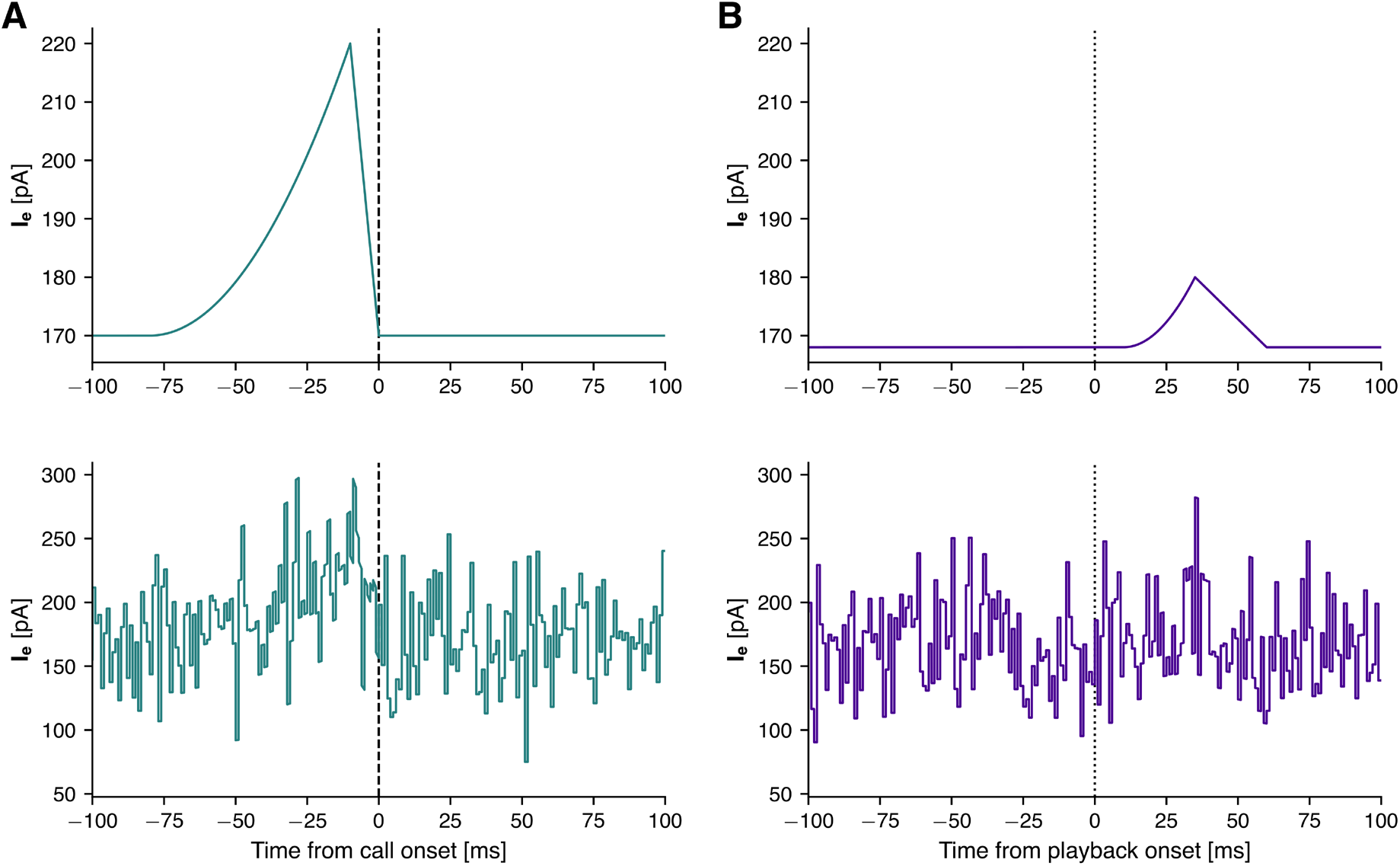
Example input currents to the upstream populations. (**A**) Input current to a neuron in the vocal-related input population, before (top) and after adding a time-varying randomized offset (bottom). (**B**) Same for the auditory-related input population.

**Figure S3 –.**
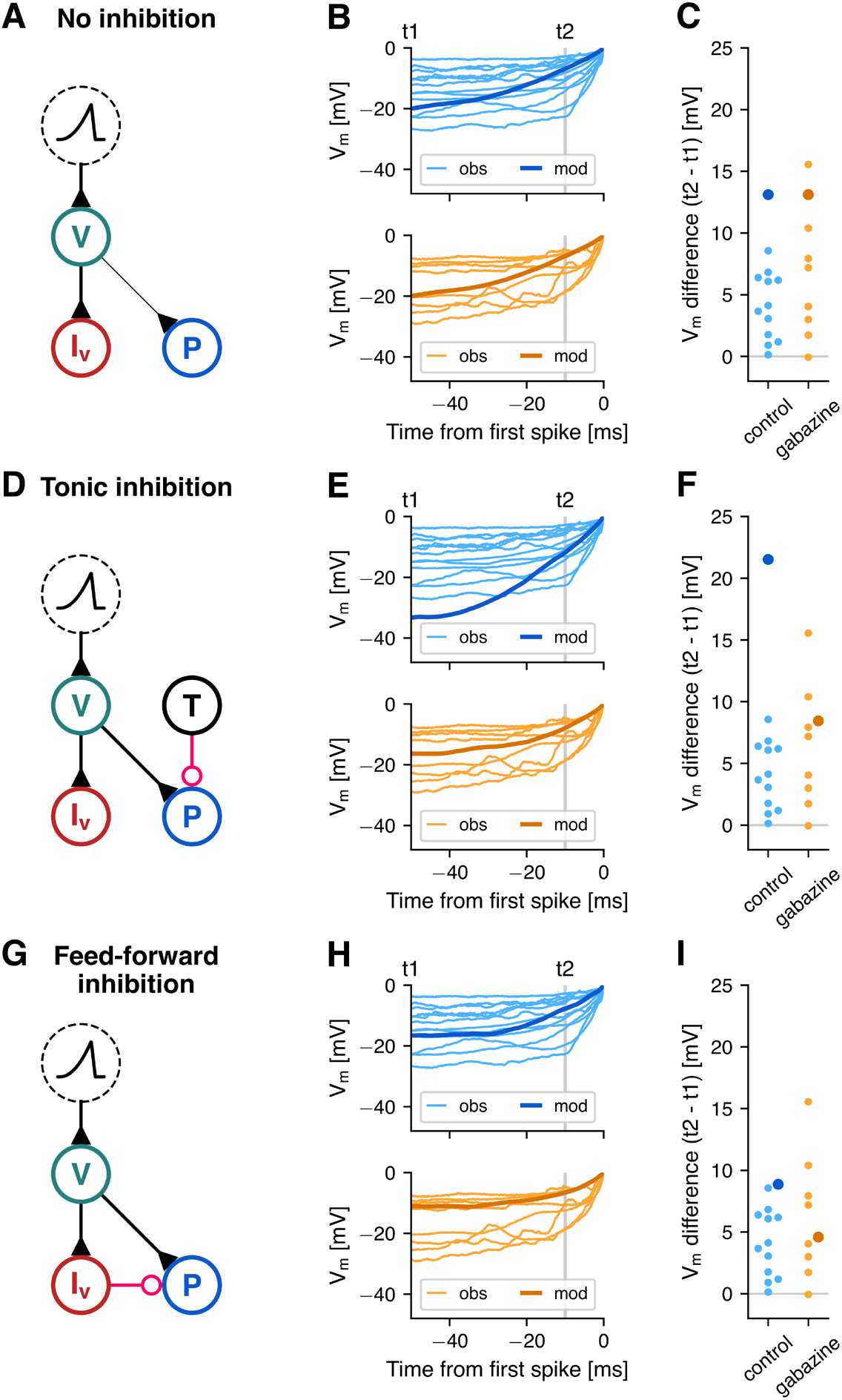
Comparison of pre-burst subthreshold potential between different modeled and observed premotor neurons. – (A, D, G) Circuit diagrams of the three model variants introduced in Figure 2. (**B, E, H**) Ramping subthreshold membrane potential of the twelve observed HVC premotor neurons that burst around call onset during the control (light blue, top) and gabazine condition (light orange, bottom), as well as the model premotor neuron at −19pA (dark blue, top) and −7pA inhibitory weights (dark orange, bottom) in the No Inhibition (B), Tonic Inhibition (E) and Feed-Forward Phasic Inhibition (H) models. Traces are aligned to the time point and the membrane potential of their first spike onset (0 ms; 0 mV). Observed traces were averaged across trials and the model traces were averaged across 100 simulations, each with different randomized amplitude offsets in the input current onto the vocal-related input neurons. (**C**, **F**, **I**) Comparison of the differences in membrane potential between 10 ms (t2) and 50 ms before burst onset (t1) for observed (small, light-colored dots) and model premotor neurons (large, dark-colored dots).

**Figure S4 –.**
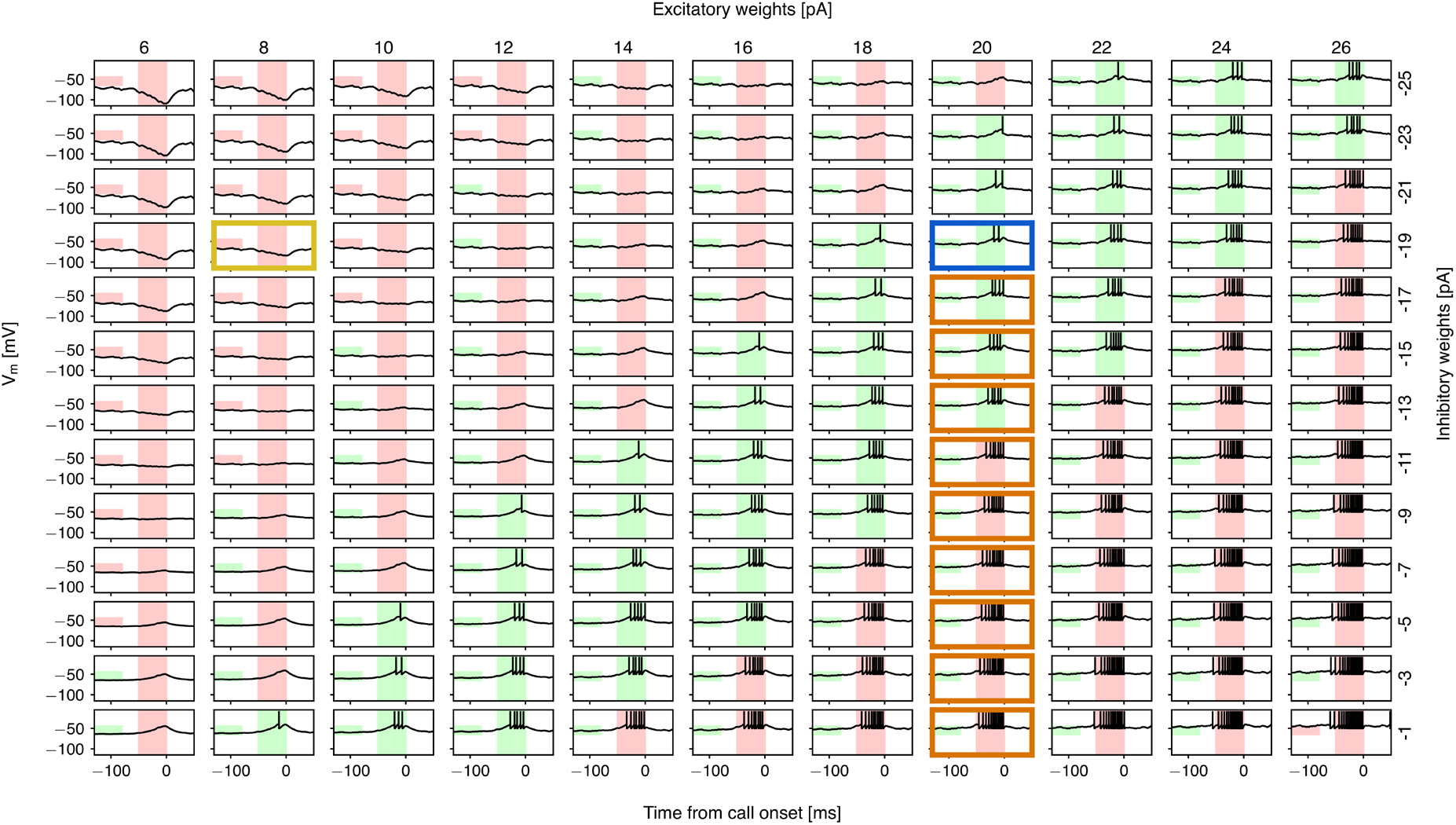
Sensitivity analysis for the Feed-forward inhibition model: synaptic weights. Membrane potential traces of the model premotor neuron in dependence on the weights of its excitatory (left to right) and inhibitory inputs (bottom to top). The colored rectangles show whether two criteria are fulfilled (green) or violated (red) in the different simulations: First (left rectangle in each panel), that the baseline membrane potential (average potential between 130 and 80 ms prior to call onset) is in the range of the recorded premotor neurons. This range is between 5 and 25 mV below spike threshold (see e.g. Figure 1E), which corresponds to −65 – −45 mV in the model. Second (right rectangle in each panel), that the neuron produces between one and six spikes during the 50ms prior to call onset, as was observed in the recorded premotor neurons. Colored frames mark the parameter combinations used in the simulations. Yellow (8 pA, −19 pA): silent premotor neuron; Blue (20 pA, −19 pA): bursting premotor neuron; Orange (20 pA, −17 – −1 pA): reduced inhibitory weights in Figure 2E–F.

**Figure S5 –.**
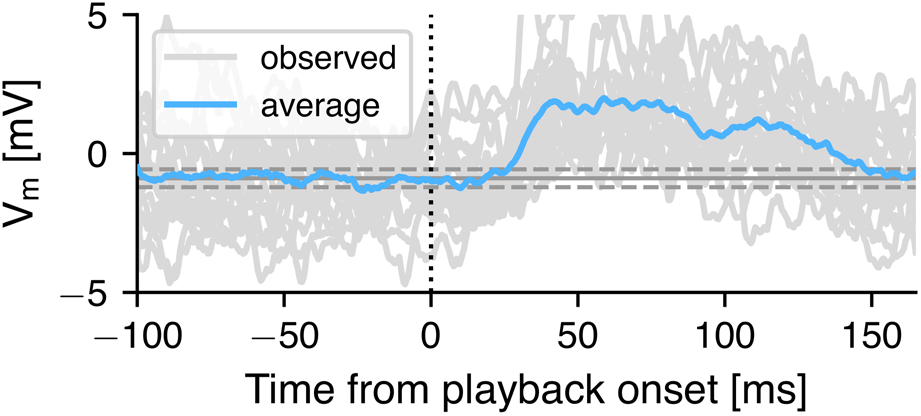
Intracellular recordings of an example premotor neuron aligned to playback onset (dotted line), which is significantly depolarized following playback onset. Horizontal lines show mean baseline potential ± 2 standard deviations (baseline: −100–0 ms).

**Figure S6 –.**
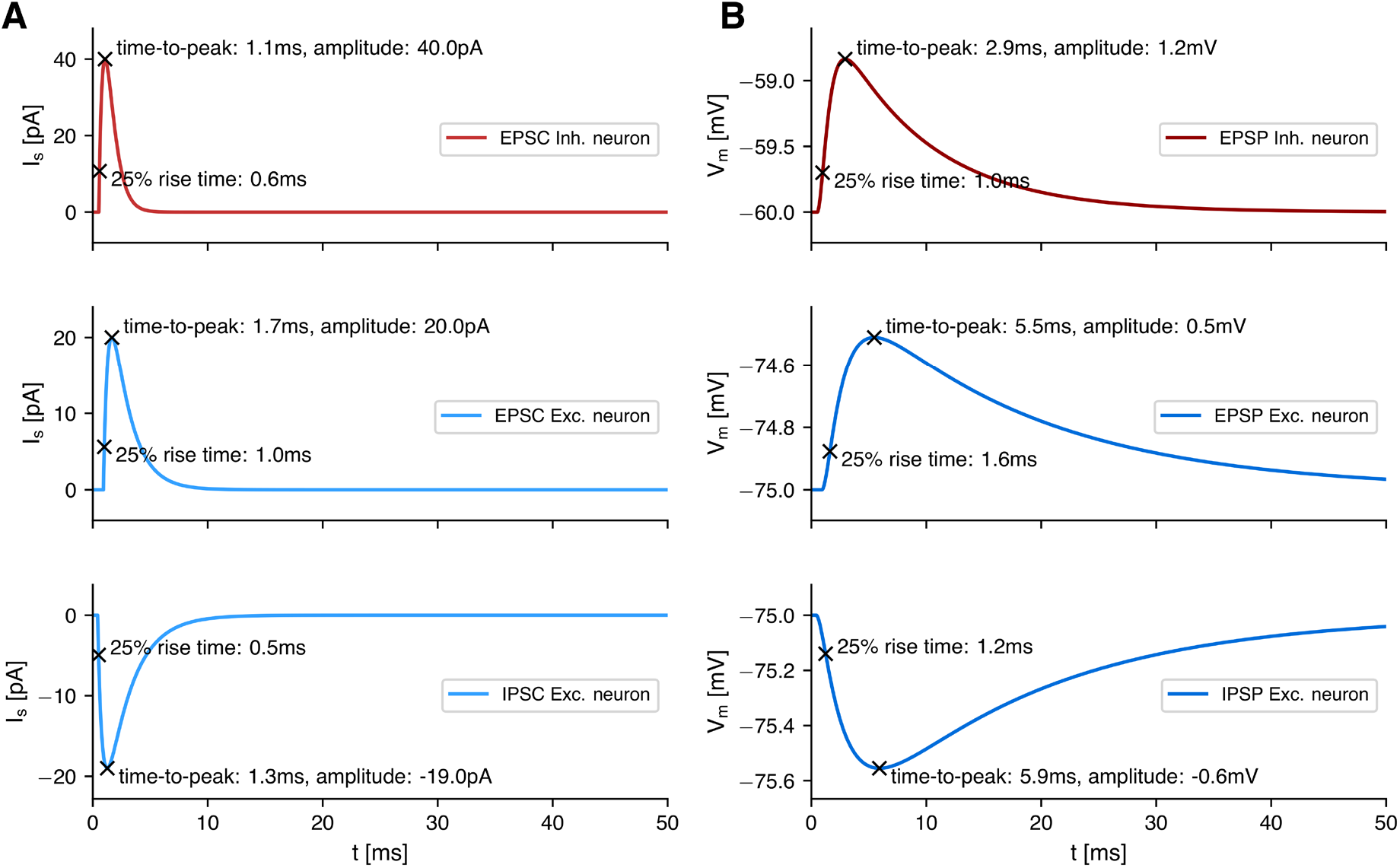
Postsynaptic currents and potentials. (**A**) Excitatory and inhibitory postsynaptic currents (EPSC/IPSC) onto an inhibitory (top) and an excitatory model neuron (middle, bottom) after a single presynaptic spike at time t=0. (**B**) Resulting postsynaptic potentials (EPSP/IPSP) in model neurons at rest.

